# A single-cell RNA-seq Training and Analysis Suite using the Galaxy Framework

**DOI:** 10.1101/2020.06.06.137570

**Authors:** Mehmet Tekman, Bérénice Batut, Alexander Ostrovsky, Christophe Antoniewski, Dave Clements, Fidel Ramirez, Graham J Etherington, Hans-Rudolf Hotz, Jelle Scholtalbers, Jonathan R Manning, Lea Bellenger, Maria A Doyle, Mohammad Heydarian, Ni Huang, Nicola Soranzo, Pablo Moreno, Stefan Mautner, Irene Papatheodorou, Anton Nekrutenko, James Taylor, Daniel Blankenberg, Rolf Backofen, Björn Grüning

## Abstract

**Background:** The vast ecosystem of single-cell RNA-seq tools has until recently been plagued by an excess of diverging analysis strategies, inconsistent file formats, and compatibility issues between different software suites. The uptake of 10x Genomics datasets has begun to calm this diversity, and the bioinformatics community leans once more towards the large computing requirements and the statistically-driven methods needed to process and understand these ever-growing datasets.

**Results:** Here we outline several Galaxy workflows and learning resources for scRNA-seq, with the aim of providing a comprehensive analysis environment paired with a thorough user learning experience that bridges the knowledge gap between the computational methods and the underlying cell biology. The Galaxy reproducible bioinformatics framework provides tools, workflows and trainings that not only enable users to perform one-click 10x preprocessing, but also empowers them to demultiplex raw sequencing from custom tagged and full-length sequencing protocols. The downstream analysis supports a wide range of high-quality interoperable suites separated into common stages of analysis: inspection, filtering, normalization, confounder removal and clustering. The teaching resources cover an assortment of different concepts from computer science to cell biology. Access to all resources is provided at the singlecell.usegalaxy.eu portal.

**Conclusions:** The reproducible and training-oriented Galaxy framework provides a sustainable HPC environment for users to run flexible analyses on both 10x and alternative platforms. The tutorials from the Galaxy Training Network along with the frequent training workshops hosted by the Galaxy Community provide a means for users to learn, publish and teach scRNA-seq analysis.

**Key Points:**

- Single-cell RNA-seq has stabilised towards 10x Genomics datasets.
- Galaxy provides rich and reproducible scRNA-seq workflows with a wide range of robust tools.
- The Galaxy Training Network provides tutorials for the processing of both 10x and non-10x datasets.

## Background

### Single-Cell RNA-seq and cellular heterogeneity

The continuing rise in single cell technologies has led to previously unprecedented levels of analysis into cell heterogeneity within tissue samples, providing new insights into developmental and differentiation pathways for a wide range of disciplines. Gene expression studies are now performed at a cellular level of resolution, which compared to bulk RNA-seq methods, allows researchers to model their tissue samples as distributions of different expressions instead of as an average.

### Pathways from Single-cell data

The various expression profiles uncovered within tissue samples infer discrete cell types which are related to one another across an “expression landscape”. The relationships between the more distinct profiles are inferred via distance-metrics or manifold learning techniques. Ultimately, the aim is to model the continuous biological process of cell differentiation from multipotent stem cells to distinct mature cell types, and infer lineage and differentiation pathways between transient cell types [1].

### Elucidating Cell Identity

Trajectory analysis which integrates the up or down regulation of significant genes along lineage branches can then be performed in order to uncover the factors and extracellular triggers that can coerce a pluripotent cell to become biased towards one cell fate outcome compared to another. This undertaking has created a new frontier of exploration in cell biology, where researchers have assembled reference maps for different cell lines for the purpose of fully recording these cell dynamics and their characteristics in which to create a global “atlas” of cells [2, 3].

### Pitfalls and Technical Challenges

#### Sequencing sensitivity and Normalization

With each new protocol comes a host of new technical problems to overcome. The first wave of software utilities to deal with the analysis of single cell datasets were statistical packages, aimed at tackling the issue of “dropout events” during sequencing, which would manifest as a high prevalence of zero-entries in over 80% of the featurecount matrix. These zeroes were problematic, since they could not be trivially ignored as their presence stated that either the cell did not produce any molecules for that transcript, or that the sequencer simply did not detect them. Normalisation techniques originally developed for bulk RNA-seq had to be adapted to accommodate for this uncertainty, and new ones were created that harnessed hurdle models, data imputation via manifold learning techniques, or by pooling subsets of cells together and building general linear models [4].

#### Improvements in sequencing

With the downstream analysis packages attempting to solve the dropouts via stochastic methods, the upstream sequencing technologies also aspired to solve the capture efficiency via new well, droplet, and flow cytometry based protocols, all of which lend importance to the process of barcoding sequencing reads.

In each protocol, cells are tagged with cell barcodes such that any reads derived from them can be unambiguously assigned to the cell of origin. The inclusion of unique molecular identifiers (UMIs) are also employed to mitigate the effects of amplification bias of transcripts within the same cell. The detection, extraction, and (de-)multiplexing of cell barcodes and UMIs is therefore one of the first hurdles researchers encounter when receiving raw `FASTQ` data from a sequencing facility.

### The Burgeoning Software Ecosystem

Since its conception, several different packages and many pipelines have been developed to assist researchers in the analysis of scRNA-seq [5, 6]. The vast majority of these packages were written for the R programming language since many of the novel normalisation methods developed to handle the dropout events depended on statistical packages that were primarily R-based [7]. Standalone analysis suites emerged as the different authors of these packages rapidly expanded their methods to encapsulate all facets of the single-cell analysis, often creating compatibility issues with previous package versions. The Bioconductor repository provided some much-needed stability in this regard by hosting stable releases, but researchers were still prone to building directly from repository sources in order to reap the benefits of new features in the upstream versions [8, 9].

#### Nonexchangeable Data Formats

Another issue was the proliferation of the many different and quickly evolving R-based file formats for processing and storing the data, such as `SingleCellExperiment` as used by the *Scater* suite, `scseq` from *RaceID*, and `SeuratObject` from *Seurat* [10, 11]. Many new packages would cater only towards one format or suite, leading to the common problem that data processed in one suite could not be reliably processed by methods in another. This incompatibility between packages fuelled a choice of one analysis suite over another, or conversely required researchers to dig deeper into the internal semantics of R `S4` objects in order to manually slot data components together [12]. These problems only accelerated the rapid development of these suites, leading to further version instability. As a result of this analysis diversity, there are many tutorials on how to perform scRNA-seq analysis each oriented around one of these pipelines [13].

#### Error propagation and Analysis Uncertainty

Different pipelines produce different results, where the stochastic nature of the analyses means that any uncertainty in a crucial quality control step upstream, such as filtering or the removal of unwanted variability, can propagate forward into the downstream sections to yield wildly different results on the same data. This uncertainty, and the statistically-driven methods to overcome them, leaves a wide knowledge gap for researchers simply trying to understand the underlying dynamics of cell identity.

### Rise of 10x Genomics

#### 10xLaunch

In 2015, 10x Genomics released their *GemCode* product, which was a droplet-seq based protocol capable of sequencing tens of thousands of cells with an average cell quality higher than other facilities [14]. This unprecedented level of throughput steadily gained traction amongst researchers and startups seeking to perform single-cell analysis, and thus 10x datasets began to prevail in the field.

#### 10xAnalysis Software

10x Genomics provided software that was able to perform much of the pre-processing, and provided feature-count matrices in a transparent HDF5-based format which provided a means of efficient matrix storage and exchange, and conclusively removed the restriction for downstream analysis modules to be written in R.

#### ScanPy, a popular alternative

The *ScanPy* suite [15], written in Python, using its own HDF5-based `AnnData` format became a valid alternative for analysing 10x datasets. The Seurat developers had similar aspirations and soon adopted the `LOOM` format, another `HDF5` variant. However, the popularity of ScanPy rose as it began to integrate the methods of other standalone packages into its codebase, making it the natural choice for users who wanted to achieve more without compromising on compatibility between different suites [9].

### Solutions in the Cloud

#### Accessible Science

As the size of datasets scaled, so did the computing resources required to analyse them, both in terms of the processing power and in storage. Galaxy is an opensource biocomputing infrastructure that exemplifies the three main tenets of science: reproducibility, peer review, and openaccess - all freely accessible within the web-browser [16]. It hosts a wide range of highly-cited bioinformatics tools with many different versions, and enables users to freely create their own workflows via a seamless drag-and-drop interface.

#### Reproducible Workflows

Galaxy can make use of *Conda* or *Containers* to setup tool environments in order to ensure that the bioinformatics tools will always be able to run, even when the library dependencies for a tool have changed, by building tools under locked version dependencies and bundling them together in a self-contained environment [17]. These environments provide a concise solution for the package version instability that plagues scRNA-seq analysis notebooks, both in terms of reproducibility and analysis flexibility. A user could keep the quality control results obtained from an older version of ScanPy, whilst running a newer ScanPy version at the clustering stage to reap the benefits of the later improvements in that algorithm. By allowing the user to select from multiple versions of the same tool, and by further permitting different versions of the tools within a workflow, Galaxy enables an unprecedented level of free-flow analysis by letting researchers pick and choose the best aspects of a tool without worrying about the underlying software libraries [18]. The burden of software incompatibility and choice of programming language that plagued the scRNA-seq analysis ecosystem before, are now completely alleviated from the user.

#### User-driven Custom Workflows

Analyses are not limited to one pipeline either, as the datasets which are passed between tools can easily be interpreted by a different tool that is capable of reading that dataset. In the case of scRNA-seq, Galaxy can convert between `CSV, MTX, LOOM` and `AnnData` formats. This interexchange of modules from different tools further extends the flexibility of the analysis by again letting the user decide which component of a tool would be best suited for a specific part of an analysis.

#### Training Resources

Galaxy also provides a wide range of learning resources, with the aim of guiding users step-by-step through an analysis, often reproducing the results of published works. The teaching and training materials are part of the Galaxy Training Network (GTN), which is a worldwide collaborative effort to produce high-quality teaching material in order to educate users in how to analyse their data, and in turn to train others of the same materials via easily deployable workshops backed by monthly stable releases of the GTN materials [19]. Training materials are provided on a wide variety of different topics, and workshops are hosted regularly, as advertised on the Galaxy Events web portal. The GTN has grown rapidly since its conception and gains new volunteers every year who each contribute and coordinate training and teaching events, maintain topic and subtopics, translate tutorials into multiple languages, and provide peer review on new material [20].

## Methods

### Stable Workflows in Galaxy

The analysis of scRNA-seq within Galaxy was a two-pronged effort concentrated on bringing high quality single-cell tools into Galaxy, and providing the necessary workflows and training to accompany them. As mentioned in the previous section, this effort needed to overcome incompatible file format issues, unstable packages due to rapid development, and needed to establish a standardised basis for the analysis.

### Tutorials

The tutorials are split into two main parts as outlined in Figure 1: first, the *pre-processing* stage which constructs a count matrix from the initial sequencing data; second, the cluster-based *downstream analysis* on the count matrix. These stages are very different from one another in terms of their information content, since the pre-processing stage requires the researcher to be more familiar with wetlab sequencing protocols than your average bioinformatician would normally know, and the downstream analysis stage requires the researcher to be familiar with statistics concepts that a wetlab scientist might not be too familiar with. The tutorials are designed to broadly appeal to both the biologist and the statistician, as well as complete beginners to the entire topic.

**Figure 1.**
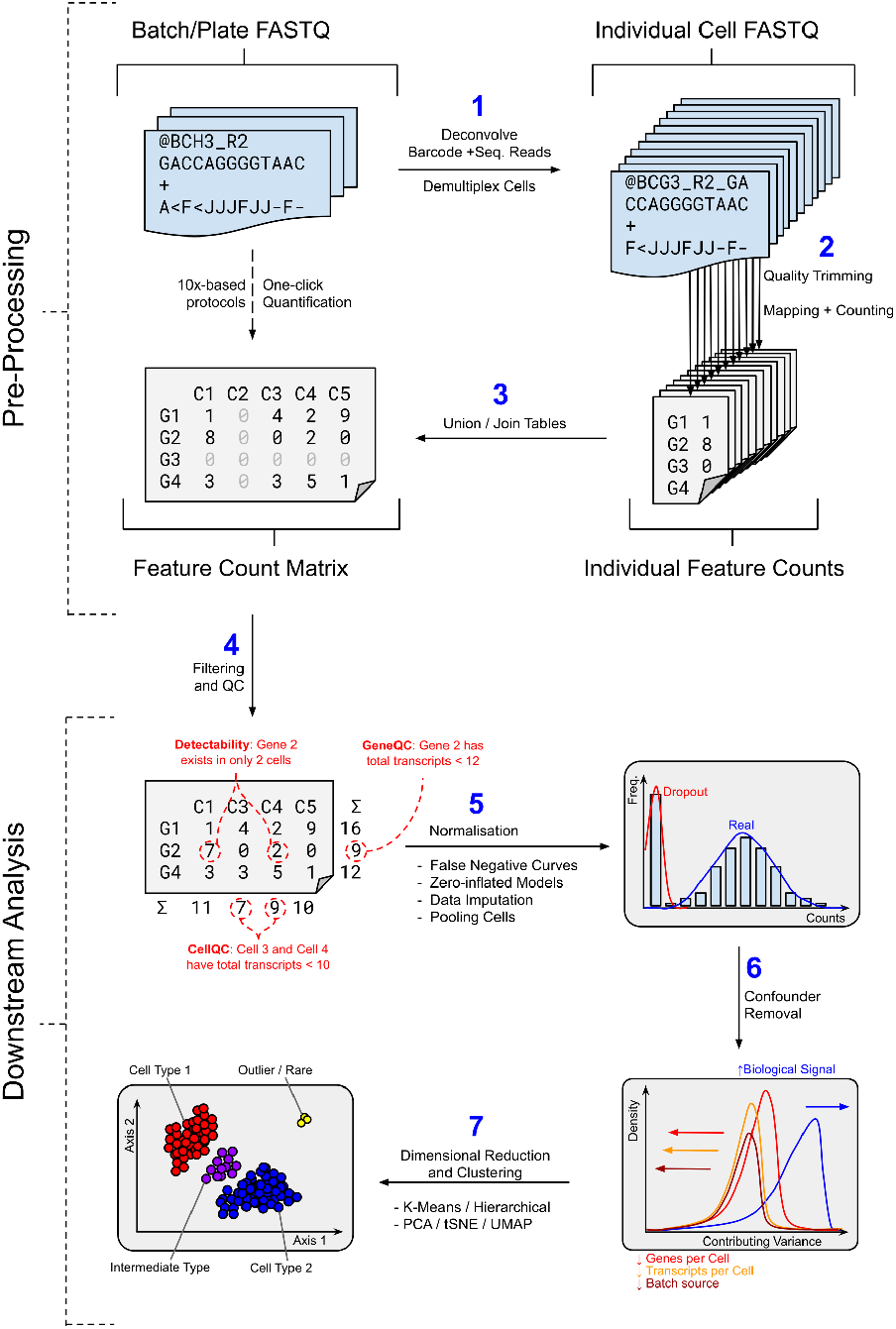
The main stages of single-cell analysis, separated broadly into the upper and lower stages of pre-processing and downstream analysis, respectively. The upper part illustrates the two main routes to generating a count-matrix from sequencing data; via one-click quantification solutions, or through manual demultiplexing. The lower part describes the four main stages required to perform cluster-based analysis from the count-matrix, through filtering, normalisation, confounder removal, and embedding.

### Pre-processing Workflows

The pre-processing scRNA-seq materials tackle the two most common use-cases that researchers will encounter when they first begin the field: processing scRNA-seq data from 10x Genomics, and processing data generated from alternative protocols. For instance, microwell-based protocols have been known to yield more features and display lower levels of dropouts compared to 10x, and so we accommodate for them by providing a more customizable path through the pre-processing stage [21].

#### Barcode Extraction

Before the era of 10x Genomics, scRNA-seq data had to be demultiplexed, mapped, and quantified. The demultiplexing stage entails an intimate knowledge of cell barcodes and Unique Molecular Identifiers (UMIs) which are protocol dependent, and expects that the bioinformatician knows exactly where and how the data was generated. One common pitfall at this very first stage is estimating how many cells to expect from the `FASTQ` input data, and this requires three crucial pieces of information: which reads contain the barcodes (or precisely, which subset of both the forward and reverse reads contains the barcodes); of these barcodes, which specific ones were actually used for the analysis; and how to resolve barcode mismatches/errors.

#### Barcode Estimation

Naive strategies involve using a known barcode template and querying against the `FASTQ` data to profile the number of reads that align to a specific barcode, often employing ‘knee’ methods to estimate this amount [22]. However, this approach is not robust, since certain cells are more likely to be over-represented compared to others, and some cell barcodes may contain more unmappable reads compared to others, meaning that the metric of higher library read depth is not necessarily correlated with a better-defined cell. Ultimately, the bioinformatician must inquire directly with the sequencing lab as to which cell barcodes were used, as these are often not specific to the protocol but to the technician who designed them, with the idea that they should not align to a specific reference genome or transcriptome.

##### One-click Pre-processing

###### Quantification with Cell Ranger

10x Genomics simplified the scRNA package ecosystem by using a language independent file format, and streamlining much of the barcode particularities with their *Cell Ranger* pipeline, allowing researchers to focus more on the internal biological variability of their datasets [23].

###### Quantification with STARsolo

The pre-processing workflow (titled “10X StarSolo Workflow”) in Galaxy uses *RNASTARsolo* utility as a drop-in replacement for *Cell Ranger*, because not only is it a feature update of the already existing *RNA STAR* tool in Galaxy, but because it boasts a ten-fold speedup in comparison to *Cell Ranger* and does not require Illumina lane-read information to perform the processing [24, 25].

###### Other Approaches

The pre-processing workflows for these “one-click” solutions consume the same datasets and yield approximately the same count matrices by following similar modes of barcode discovery and quantification. Within Galaxy, there is also *Alevin* (paired with *Salmon*) and *scPipe* which can both also perform the necessary demultiplexing, (alignment-free) mapping, and quantification stages in a single step [26, 27, 28].

##### Flexible Pre-processing

###### CELSeq2 Barcoding

The custom pre-processing workflow (titled “CELSeq2: Single Batch mm10”) is modelled after the CEL-seq2 protocol using the barcoding strategies of the Freiburg Max-Planck Institute laboratory as its main template, but the workflow is actually flexible to accommodate any droplet or well-based protocol such as SMART-seq2, and Drop-seq [29].

###### Manual Demultiplexing and Quantification

The training picto-graphically guides users through the concepts of extracting cell barcodes from the protocol, explains the significance of UMIs in the process of read deduplication with illustrative examples, and instructs the user in the process of performing further quality controls on their data during the post-mapping process via *RNA STAR* and other tools that are native to Galaxy.

###### Training the User

At each stage, the user’s knowledge is queried via question prompts and expandable answer box dialogs, as well as helpful hints for future processing in comment boxes, all written in the transparent Markdown specification developed for contributing to the GTN.

### Downstream Workflows

#### Common Stages of Analysis

The downstream modules are defined by the five main stages of downstream scRNA-seq analysis: filtering, normalisation, confounder removal, clustering, and trajectory inference. There are three workflows to aid in this process (two of which are shown in Figure 2), each sporting a different well-established scRNA-seq pipeline tool.

**Figure 2.**
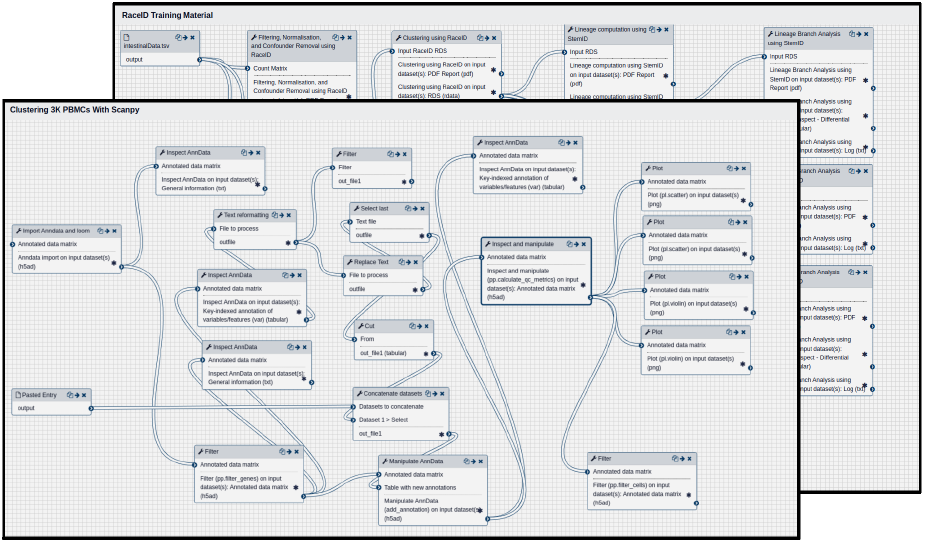
Downstream analysis workflows as shown in the Galaxy Workflow Editor for (top) RaceID and (bottom) ScanPy, each displaying modules symbolizing the five main stages of analysis.

#### Quality Control with Scater

The Scater pipeline follows a visualise-filter-visualise paradigm which provides an intuitive means to perform quality control on a count matrix by use of repeated incremental changes on a dataset through the use of PCA and library size based metrics [30]. Once this pre-analysis stage is complete, the full downstream analysis (comprising the five stages mentioned above) can be performed by workflows based on the following suites: RaceID and ScanPy.

#### Downstream Analysis with the RaceID Suite

RaceID was developed initially to analyse rare cell transcriptomes whilst being robust against noise, and thus is ideal for working with smaller datasets in the range of 300 to 1000 cells. Due to its complex cell lineage and fate predictions models, it can also be used on larger datasets with some scaling costs.

#### Downstream Analysis with the ScanPy Suite

ScanPy was developed as the Python alternative to the innumerable R-based packages for scRNA-seq which was the dominant language for such analyses, and it was one of the first packages with native 10x genomics support. Since then it has grown substantially, and has been re-implementing much of the newer R-based methods released in BioConductor as “recipe” modules, thereby providing a single source to perform many different types of the same analysis.

The workflows derived from both these suites emulate the five main stages of analysis mentioned previously, where filtering, normalisation, and confounder removal are typically separated into distinct stages, though sometimes merged into one step depending on the tool.

##### Filtering

###### Cell and Gene Removal

During the filtering stage, the initial count matrix removes low-quality or unwanted cells using commonly used parameters such as minimum gene detection sensitivity and minimum library size, and low-quality genes are also removed under similar metrics, where the minimum number of cells for a gene to be included is decided. The Scater pre-analysis workflow can also be used here to provide a PCA-based method of feature selection so that only the highly variable genes are left in the analysis.

###### Disadvantages of Filtering

There is always the danger of over-filtering a dataset, whereby setting overzealous lower-bound thresholds on gene variability, can have the undesired effect of removing essential housekeeping genes. These relatively uniformly expressed genes are often required for setting a baseline to which the more desired differentially expressed genes can be selected from. It is therefore important that the user first performs a naive analysis and only later refine their filtering thresholds to boost the biological signal.

##### Normalisation

###### Library Size Normalisation

The normalisation step aims to remove any technical factors that are not relevant to the analysis, such as the library size, where cells sharing the same identity are likely to differ from one another more by the number of transcripts they exhibit, than due to more relevant biological factors.

###### Intrinsic Cell Factors

The first and foremost is cell capture efficiency, where different cells produce more or less transcripts based on the amplification and coverage conditions they are sequenced in. The second is the presence of dropout events which manifest as a prevalence of “zeroes” in the final count matrix. Whether a “zero” is imputable to the lack of detection of an existing molecule or to the absence of the molecule in the cell is uncertain. This uncertainty alone has led to a wide selection of different normalisation techniques that try to model this expression either via hurdle models, or imputing the data via manifold learning techniques, or working around the issue by pooling subsets of cells together [31].

In this regard, both the RaceID and ScanPy workflows offer many different normalisation techniques, and users are encouraged to take advantage of the branching workflow model of Galaxy to explore all possible options.

##### Confounder Removal

###### Regression of Cell Cycle Effects

Other sources of variability stem from unwanted biological contributions known as confounder effects, such as cell cycle effects and transcription. Depending on what stage of the cell cycle a cell was sequenced at, two cells of the same type might cluster differently because one might have more transcripts due to it being in the M-phase of the cell cycle. Library sizes notwithstanding, it is the variability in specific cell cycle genes that can be the main driving factor in the overall variability. Thankfully, these effects are easy to regress out, and we replicate an entire standalone ScanPy workflow dedicated to detecting and visualising the effects based on the original notebook [32].

###### Transcriptional Bursting

The transcription effects are harder to model, as these are semi-stochastic and are as of yet still not well understood. In bulk RNA-seq the expression of genes undergoing transcription are averaged to give “high” or “low” signals producing a global effect that gives the false impression that transcription is a continuous process. The reality is more complex, where cells undergo transcription in “bursts” of activity followed by periods of no activity, at irregular intervals [33]. At the bulk level these discrete processes are smoothed to give a continuous effect, but at the cell level it could mean that even two directly adjacent cells of the same type normalised to the same number of transcripts can still have different levels of expression for a gene due to this process. This is not something that can be countered for, but it does educate the users in which factors they can or cannot control in an analysis, and how much variability they can expect to see.

##### Clustering and Projection

###### Dimension Reduction and Clustering

Once a user has obtained a count matrix they are confident with, they are then guided through the process of dimension reduction (with choice of different distance metrics), choosing a suitable low-dimensional embedding, and performing clustering through commonly-used techniques such as k-means, hierarchical, and neighbourhood community detection. These complex techniques are illustrated in layman’s terms through the use of helpful images and community examples. For example, the GTN ScanPy tutorial explains the Louvain clustering approach[34] via a standalone slide deck to assist in the workflow [35].

###### Commonly-used Embeddings

The clustering and the cluster inspection stages are notably separated into distinct utilities here, with the understanding that the same initial clustering can appear dissimilar under different projections, e.g. t-distributed Stochastic Network Embedding (tSNE) against Uniform Manifold Approximation and Projection (UMAP) [36, 37]. Ultimately the user is encouraged to play with the plotting parameters to yield the best looking clusters.

###### Static Plots or Interactive Environments

Cluster inspection tools are available that allow users to easily generate static plots tailored to pipeline-specific information as originally defined by the software package authors. However, the `AnnData` and `LOOM` specifications store this map projection data separately, enabling the use of a plethora of possible plotting tools, including HTML5-based interactive visualisations, such as *cellx-gene*[38], that permit on-demand querying and rendering of individual cell features without having to generate static images. A collection of these Galaxy interactive tools can be accessed at the website live.usegalaxy.eu. Though these tools are excellent at dynamically displaying map projections, especially 3-dimensional ones, further computation must be performed to complete a full pseudotime analysis.

##### Pseudotime Trajectory Analysis

###### Inferring Developmental Pathways

The cell pseudotime series analysis is often referred to as the trajectory inference stage, since cells are ordered along a trajectory to reflect the continuous changes of gene expression along a developmental pathway under the assumption that the cells are transitioning from one pluripotent type to another less-potent type.

###### Pseudotime Techniques

For the trajectory inference stage, there is the Partition-based Graph Abstraction (PAGA) technique championed by ScanPy [39], and there is also the FateID and StemID packages for the RaceID workflow [40]. The former provides a level of graph abstraction to the datasets in order to infer a community graph structure, which it can use to learn the shape of the data and infer pathways between neighbourhoods. The latter is more intuitive, in that it constructs a minimal spanning tree of related clusters that infer lineage, and cell fate decisions that can be explored by querying branches in the tree, as a function of the genes which are up or down regulated along the currently explored pathway. The statistical strength and significance of each pathway guides the user along more valid trajectories that would more accurately reflect the biological variation occurring within transitioning cells.

##### Sharing Reference Maps

The insights and novel cell types discovered in these analyses can also be integrated into the Human Cell Atlas portal [41], which is an initiative that aims to classify unique or rare cell types as well as their transitive properties in order to build a comprehensive map of cells that can be used to investigate the various differentiation pathways of multipotent stem cells in the human body.

### Galaxy Training Network

#### Tutorial Hierarchy

Tutorials in the GTN are grouped by topic, e.g. Variant Calling, Transcriptomics, Assembly, etc. These tutorials can also declare prerequisites, so that users can review required concepts from previous tutorials, e.g. quality control checks from bulk RNA-seq still being used in scRNA-seq. Not only does this allow users to derive a clear route through the range of training materials, but it also empowers them to choose their own learning path through the network of topics. In particular for scRNA-seq, users can start their training from pre-processing tutorials and continue till downstream analysis.

#### Tutorial Structure

Tutorials usually consist of a hands-on workflow that guides the user through an analysis with Galaxy utilising a step-by-step approach, and is often accompanied with a slide deck that either serves to explain standalone concepts more concisely, or is used during workshops and trainings as a way to introduce the user to the topic. In an effort to maintain reproducibility in science, all tutorials require example workflows, and all materials needed to run the workflows and tutorials are hosted for free with open access at Zenodo with a permanent DOI tag.

#### User-driven Contribution

The user contributions are the heart of the GTN community, and options are given to appeal to different levels of contribution. At the casual level, each tutorial has at the bottom an anonymous feedback form that rates the quality of the tutorial and asks for hints on what could be improved, which the tutorial authors can then act on. At the more eager level, users can contribute directly to the material hosted at the GitHub repository using the approachable GTN Markdown format, which further empowers contributors to not only adapt existing material, but to also write tutorials in their own specialist topics. The GitHub code reviews paired with the plaintext GTN Markdown format, facilitate easy peer-review of tutorial topics by using standard diff utilities.

### Galaxy Subdomains and Environments

#### Subdomains Encapsulate Relevant Tools

The Galaxy tools and the GTN are further tied together by Galaxy subdomains, that better serve the various topics within their own self-contained environments. These complement the training materials by providing only the necessary Galaxy tools in order to not trouble the user with unrelated tools that might not be so relevant to the material, e.g. Variant Analysis tools are not included in an scRNA-seq environment. This also has the benefit that smaller more specialised Galaxy instances can be packaged and deployed, avoiding the overhead of presenting the entire Galaxy tool repertoire.

#### Single Cell and Human Cell Atlas

In this light, the singlecell. usegalaxy.eu subdomain hosts the entirety of the single-cell materials, tools, workflows, and single-cell related events. A table containing the full list of tools in the subdomain, as well as their application to the previously mentioned stages of scRNA-seq analysis is given in Supplemental Table 1. Human Cell Atlas community members, led by the European Bioinformatics Institute and the Wellcome Sanger Institute have their own subdomain at humancellatlas.usegalaxy.eu [42], providing access to widely applicable tools including ScanPy, Seurat and *Monocle3* [43], but also specialist tools such as those for cell type prediction (including *scmap* [44], *scPred* [45] and *Garnett* [46]).

#### Analysis in Galaxy workflows

The tools outlined in the *Downstream Workflows* subsection expose the full set of parameters of their underlying program suites, in order to serve the same level of customisation that the users would expect when running a notebook-based analysis. This suits the needs of most researchers, but some are more used to processing the data directly in a language-driven notebook environment.

#### Galaxy Interactive Environments (GIE)

For the more computer programming-oriented users, Galaxy hosts interactive environments at live.usegalaxy.eu which allows users to spin up their own Jupyter [47] or RStudio [48] notebooks whilst harnessing the same cloud compute infrastructure. Here, users can import their Galaxy datasets, process them in their own desired manner, and export them back into their histories in a similar way to how datasets are treated in workflows.

#### List of GIEs

In addition to interactive notebooks, the GIE also boasts a selection of other interactive tools such as the previously mentioned *cellxgene* featured in Figure 3, as well as *SPARQL* a query language interface, *BAM/VCF IOBIO* a file format analysis viewer [49], *EtherCalc* a web spreadsheet [50], *PHINCH* a metagenomes visualiser [51], *Wallace* a species modelling platform [52], *WILSON* an omics visualiser [53], *IDE* for materials science, *Panoply* a netCDF viewer [54], *HiGlass* a Hi-C data visualiser [55], and even an XFCE Virtual Desktop environment [56].

**Figure 3.**
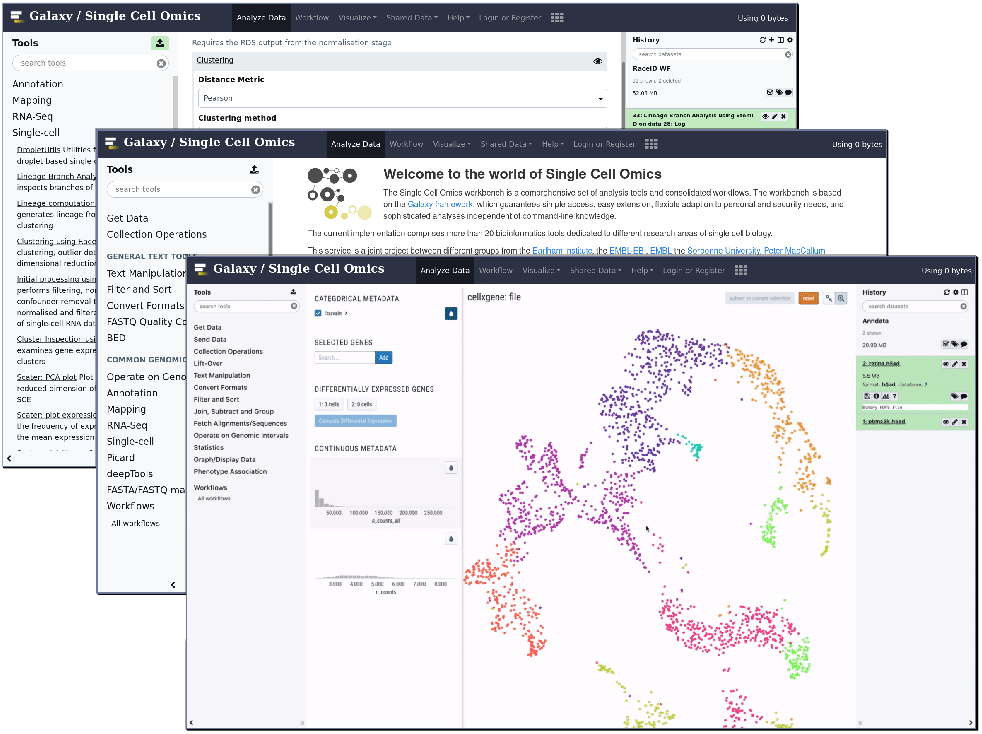
Galaxy Training Network hosting a comprehensive suite of tools, trainings, and workflows to perform scRNA-seq analysis.

## Discussion

### Growth of scRNA Training Materials

The single-cell materials on the GTN are growing substantially every year, with at first only one pre-processing tutorial in 2018, one downstream tutorial at the start of 2019, and at the current time of writing three pre-processing tutorials and three downstream analysis workflows, further accompanied by slide decks and interactive visualisations. Single-cell Galaxy workshops based on these materials have been given at the Single-Cell RNAseq Training Course 2018 at the Earlham Institute, the 2019 Galaxy Community Conference (GCC2019), within the Freiburg MeInBio consortium, and at the Association of Biomolecular Resource Facilities (ABRF). The trainings also lend themselves seamlessly to online Webinars which have proved useful during the COVID-19 lockdown period.

### Reproducible Cloud-based Analysis

The advent of scRNA-seq analysis within the Galaxy framework re-echoes the efforts to standardise the analysis of scRNA-seq with the promise of presenting reproducible research. The burden of computation on the ever-growing size of the datasets is shifted to the cloud computing resources, and as scRNA sequencing technology scales, more researchers are likely to migrate towards cloud-based solutions in order to reap the benefits of superior computing abilities and storage capabilities. Ultimately, the Galaxy framework abstracts the user from the many nontrivial technicalities of the analysis, and exposes them to a legible interface of tools that they can pick and choose from.

### Longevity and Accessibility

The community regularly comes together during scheduled code festivals (CoFests) or hackathons to review, contribute, and actively maintain the training materials. The number of community contributions have steadily increased over the last four years [16], and this growing trend ensures that the Galaxy resources will stay current and adapt to changes in scRNA sequencing technology and analysis methods if necessary. The GTN also makes use of language translation tools to provide international interpretations of the training materials in order to reach a wider more internationally diverse audience.

### Future of scRNA-seq in Galaxy

The capacity for growth of scRNA-seq in Galaxy is limitless, with the continuing acquisition of new single-cell tools being incorporated into Galaxy workflows, and the expanding GTN community bringing more expert-level contributions to the training material. The vestiges of incompatible libraries and in-exchangeable file formats are unburdened from the user as the epoch of web-based tools and strong biocomputing frameworks become more dominant. From the first data upload to the final finishing touches of a customized workflow, the single cell Galaxy portal upholds the ideals of open science by supporting the user all the way from the initial training to the final publication, where they can export and share their results with a single click.

## Availability of source code and requirements (optional, if code is present)

Lists the following:

- Project name: Single-Cell RNA-seq Analysis in Galaxy
- Project home page: singlecell.usegalaxy.eu
- Operating system(s): Web-based, Platform independent
- License: GNU GPL v3 Any restrictions to use by non-academics: e.g. licence needed

## Availability of supporting data and materials

All datasets used in the GTN are independently hosted at Zenodo and are easily findable under the tag “Galaxy Training Network”, as well as being directly hosted within the Galaxy Data Libraries on the UseGalaxy.eu server.

The tool wrappers which serve as the functional components of the many different single-cell analysis tools are hosted at the *GitHub Tools-IUC* repository, as well as at the Galaxy Toolshed under the category of “Transcriptomics”.

## Declarations

### List of abbreviations

DOI: Digital Object Identifier
GTN: Galaxy Training Network
HDF5: Hierarchical Data Format 5
HPC: High Performance Computing
PAGA: Partition-based Graph Abstraction
PCA: Principal Component Analysis
scRNA: Single-Cell RNA
tSNE: t-distributed Stochastic Network Embeddings
UMAP: Uniform Manifold Approximation and Projection
UMI: Unique Molecular Identifier

## Consent for publication

Not applicable.

## Competing Interests

The author(s) declare that they have no competing interests.

## Funding

Funded by the Deutsche Forschungsgemeinschaft (DFG, German Research Foundation) 22977937/GRK2344 and BA2168/3-3, the BBSRC Core Strategic Programme Grants BBS/E/T/000PR9814, BBS/E/T/000PR9817, BBS/E/T/000PR9818, and BBS/E/T/000PR9819, Core Capability Grant BBS/E/T/000PR9816 at the Earlham Institute, and the National Institutes of Health grant U41HG006620.

The European Galaxy project is in part funded by Collaborative Research Centre 992 Medical Epigenetics (DFG grant SFB 992/1 2012) and German Federal Ministry of Education and Research (BMBF grants 031 A538A/A538C RBC, 031L0101B/031L0101C de.NBI-epi).

## Author’s Contributions

B.G. conceived the project, and created the singlecell subdomain. M.T. and B.B. wrapped RaceID and ScanPy respectively, and created all initial workflows and trainings. A.O., B.G., C.A., D.C., D.B., G.J.E, H.R.H., J.R.M., L.B., M.D., M.H., N.H., F.R., J.S., N.S., P.M. and S.M. developed tools or made functional contributions to the tools and training materials. D.B., I.P., A.N., J.T. and R.B. supported the development of the project. M.T. wrote the original draft manuscript. All authors have read, made suggestions, and ultimately approved the final manuscript.

## Acknowledgements

We thank the bioinformatics group at the University of Freiburg for the development and hosting of the European Galaxy server, Monika Degen-Hellmuth at the Backofen Lab for her assistance in the organization of the project, the Institut Français de Bioinformatique (IFB) for its support of the ARTbio team, Charles Girardot at EMBL Heidelberg for his useful feedback, and we also thank the worldwide contributions from users and developers towards the Galaxy Project and all upstream authors and contributors of the software ecosystem that we use and rely on.

## Notes

https://singlecell.usegalaxy.eu/

### Competing Interest Statement

The authors have declared no competing interest.

### Summary of Updates

Added reviewer suggested edits; abstract wording change, minor headers changed in main text, and grammar.

